# Hematopoietic stem cell-derived conditioned media induces excessive mitochondrial fission via Drp-1 to target colorectal cancer

**DOI:** 10.1101/2025.09.10.675059

**Authors:** Sumit Mallick, Akhila Balakrishna Rai, Vanya Kadla Narayana, Sudheer Shenoy P, Thottethodi Subrahmanya Keshava Prasad, Bipasha Bose, Siddhartha Biswas

## Abstract

Mitochondria, often referred to as the “powerhouses of the cell,” are particularly crucial in cancer cells due to their high energy demands. Mitochondrial fusion-fission dynamics play a critical role in regulating signaling pathways and metabolic activities in colorectal cancer (CRC) cells. Increased mitochondrial fission drives metabolic reprogramming, enabling CRCs to proliferate, metastasize, and resist chemotherapy. Paradoxically, excessive fission induces mitochondria-mediated apoptosis. Our previous studies have shown that hematopoietic stem cell-derived conditioned media (CM) modulate the apoptosis pathway and mitochondrial bioenergetics of cancer stem cells by altering the cancer microenvironment. In this study, We found that HSCs-CM facilitates excessive fission in colorectal cancer cells by modulating Drp-1 and concurrently activating the PINK1–Parkin mitophagy pathway and intrinsic apoptosis, leading to loss of viability of these cells. Moreover, proteomics data showed that HSCs-CM dysregulated the electron transport chain complexes, with an exceptionally high degree of dysregulation of complexes III and IV. Metabolomics revealed dysregulation of critical metabolites, RNA sequencing revealed dysregulation of transcripts, and proteomics revealed dysregulation of proteins, involved in mitochondrial bioenergetics and the autophagy pathway in CRCs treated with CM. Taken together, our studies reveal the therapeutic potential of HSC-conditioned media for treating colorectal cancer.

**Highlights:**

- Hematopoietic stem cell-derived conditioned media (HSC-CM) induces excessive mitochondrial fission in colorectal cancer (CRC) cells by upregulating Drp-1, thereby activating the intrinsic apoptotic pathway.
- Excessive fission and bioenergetic dysfunction caused by HSC-CM result in loss of mitochondrial membrane potential and elevated reactive oxygen species production.
- HSC-CM severely disrupts the electron transport chain in CRC cells, precipitating an energy crisis and engaging the PINK1-mediated mitophagy pathway.

**Summary:** Hematopoietic stem cell-derived conditioned media (HSC-CM) compromises mitochondrial dynamics by inducing excessive fission in colorectal cancer (CRC) cells through upregulation of Drp-1 and its associated protein complex. The resulting hyperfission leads to severe mitochondrial dysfunction, characterised by a loss of mitochondrial membrane potential, decreased ATP production, and disrupted electron transport chain complexes. This bioenergetic crisis and overproduction of reactive oxygen species ultimately trigger intrinsic apoptosis and engage the mitophagy pathway, leading to loss of CRC cell viability.

## 1. Introduction

Colorectal cancer (CRC) is the third most prevalent cancer worldwide, with approximately 2 × 10 new diagnoses and 1 × 10 deaths reported each year. The overall 5-year survival rate is around 64%, but drops to as low as 15% in metastatic disease (mCRC) as a consequence of therapy failure and metabolic reprogramming, characterised by enhanced glycolysis and decreased oxidative phosphorylation (OXPHOS) [1–3]. Mitochondrial dynamics, comprising fission, fusion and mitophagy, are central to cancer progression and particularly relevant in mCRC, where survival and dissemination depend on adaptive metabolic plasticity. Fusion, mediated by OPA1 and MFN1/2, preserves mitochondrial function and sustains OXPHOS, a process required for energy-intensive activities such as metastasis and invasion. In contrast, fission, driven by the dynamin-related GTPase Drp-1 (encoded by DNM1L), facilitates mitochondrial fragmentation, enabling efficient organelle distribution during cell division and adaptation to hypoxic or nutrient-poor microenvironments commonly found in metastatic niches [4–6]. Dysregulated fission, however, promotes excessive mitochondrial fragmentation that culminates in bioenergetic stress, reactive oxygen species (ROS) overproduction and engagement of mitophagy-mediated apoptosis. In aggressive CRC subtypes with poor prognosis, hyperactive fission also supports glycolysis, accelerates cell-cycle entry and helps tumour cells evade mitochondrial apoptosis by sequestering damaged mitochondria via fission, thereby escaping mitophagy-dependent clearance.

Pharmacological inhibition of Drp-1 (e.g. with Mdivi-1) has shown promise in preclinical models by reversing fission, restraining metastasis and enhancing chemotherapy sensitivity [7]. Conversely, hyperactivation of Drp-1, as we observe in CRC cells exposed to hematopoietic stem cell-derived conditioned media (HSC-CM), leads to pathological mitochondrial hyperfission, irreparable metabolic derangement and apoptosis [8,9]. This dual-edged regulation underscores the plasticity of Drp-1: inhibition can blunt pro-metastatic fission, whereas hyperactivation can drive lethal fragmentation in drug-resistant mCRC cells [10]. Hematopoietic stem cells (HSCs) are best known for sustaining haematopoiesis and contributing to tissue repair, but the various cytokines and secretory molecules present in their conditioned media (HSC-CM) have not been investigated as modulators of CRC mitochondria for therapeutic purposes. In our previous work, we showed that HSC-CM is enriched in metabolites with anti-tumour potential against cancer and cancer stem cells [11–12].

Whereas our earlier studies primarily focused on apoptosis induction in cancer stem cells through disruption of HSP90 and the 26S proteasome system, the present study introduces a distinct and original mechanistic perspective. Here, for the first time, we demonstrate that HSC-CM specifically targets mitochondrial dynamics in CRC cells, shifting the balance from fusion to excessive fission, precipitating bioenergetic collapse and impairing OXPHOS. We identify the Drp-1/Fis-1 signalling axis as the principal extracellular target, indicating that the HSC secretome is capable of more than maintaining protein-quality control — it can precisely target mitochondrial structure and metabolic flexibility in aggressive CRC. Our findings establish that HSC-CM induces excessive fission leading to apoptosis, and define a metabolic vulnerability that can be therapeutically exploited in CRC.

## 2. Materials and Methods

### 2.1 Cell culture

Human colorectal cancer cells (HCT-116) and the human myeloid leukaemia line HL-60 were procured from the National Centre for Cell Science (NCCS), Pune, India. All cell lines were authenticated by the supplier using short tandem repeat (STR) profiling and were certified mycoplasma-free. HCT-116 cells were cultured in DMEM supplemented with 10% foetal bovine serum (FBS) and were trypsinised for downstream experiments at approximately 70% confluency. HL-60 cells were grown in growth-factor-enriched media (SCF; 50 ng/mL, FLT3; 20 ng/mL, IL 6; 50 ng/mL, TPO; 10 ng/mL) (Pepro Tech, ThermoFisher) & (Extended File, Table 3) [11–12]. At confluency, HL-60 cells were collected, stained with hematopoietic stem-cell markers (CD34 /CD45) and sorted by fluorescence-activated cell sorting (FACS). The sorted CD34 cells were used for downstream conditioned-media preparation (Extended File, Table 5).

### 2.2 Conditioned-media preparation

Conditioned media (CM) were prepared as described previously [11–12]. Briefly, sorted HSCs were cultured at a density of 5 × 10 cells/mL in growth-factor-enriched media for 48 h. This medium comprised serum-free basal medium supplemented with stem cell factor (SCF), FLT3 ligand, interleukin-6 (IL-6) and thrombopoietin (TPO); the complete formulation and concentrations are given in Extended File, Table 3. The supernatant was then sequentially centrifuged at 1,000 × g for 5 min, 4,000 × g for 15 min and 6,000 × g for 10 min, all at 4 °C. The final supernatant was filtered through a 0.22-µm filter to remove residual debris, aliquoted, and stored at –80 °C until further use. Throughout the study the control was the identical growth-factor-enriched medium that was not conditioned by cells, but was subjected to the same 48-h incubation, sequential centrifugation, 0.22-µm filtration, –80 °C storage and thawing, and 1:1 dilution into HCT-116 growth medium, so that conditioning by the sorted CD34 population was the only variable between groups.

### 2.3 Metabolomics

For untargeted metabolomic profiling, HCT-116 cells were treated with HSC-CM for 48 h. The medium was replaced 2 h before extraction; 80% chilled methanol was added, and the cells were incubated at –80 °C for 20 min. The cells were then scraped, centrifuged, and the resulting supernatant was supplemented with an additional methanol wash, centrifuged, dried in a SpeedVac, and resuspended in 0.1% formic acid. Samples were analysed on a Q Exactive Orbitrap mass spectrometer with full MS scans acquired at 140,000 resolution (100–700 m/z), AGC target 1 × 10 and maximum IT 50 ms. Precursors were fragmented using HCD (NCE 35%), and fragment ions were detected at 35,000 resolution (AGC 1 × 10, dynamic exclusion 10 s, loop count 5). Metabolite identification was carried out using Compound Discoverer v3.2 (Thermo Fisher Scientific) [11].

### 2.4 Proteomic analysis

HSC-CM-treated HCT-116 cells were lysed in SDS buffer containing protease inhibitors, sonicated and centrifuged. Protein concentration was determined by BCA assay; 30 µg of protein was reduced, alkylated, filtered through a 30-kDa column, and digested with trypsin at 37 °C for 16 h. The resulting peptides were desalted on C18 tips, dried, and stored prior to LC-MS/MS analysis. Peptides were analysed on an EASY-nLC 1200 coupled to an Orbitrap Fusion Tribrid mass spectrometer with full MS scans at 120,000 resolution (350–1,500 m/z) and HCD fragmentation (NCE 35%), followed by MS2 at 30,000 resolution, in technical triplicates. Data were processed in DIA-NN v1.8.1 in library-free mode against the UniProt human proteome (carbamidomethylation as fixed; methionine oxidation and N-terminal acetylation as variable modifications), with a 1% FDR cut-off, isoform-level protein inference and median normalisation [12].

### 2.5 Immunocytochemistry

HCT-116 cells were cultured with 50% HSC-CM in growth medium (1:1 v/v) for 48 h, fixed in 4% paraformaldehyde, permeabilised, and stained with primary and secondary antibodies as listed in Extended File, Table 2. Hoechst 33342 was used to visualise nuclei. Confocal images were acquired on a Zeiss LSM 880 microscope at 63× magnification. Fluorescence intensity was quantified using ImageJ, and nuclear localisation of proteins was assessed by Pearson’s correlation coefficient. Six independent fields per condition were quantified and presented as bar graphs.

### 2.6 JC-1 staining

HCT-116 cells (2 × 10 cells/well) were seeded in 6-well plates. After 6 h, cells were treated with a 1:1 mixture of HSC-CM and growth medium for 48 h. Mitochondrial membrane potential was assessed by JC-1 staining (400 nM; Invitrogen, T3168), where red fluorescence indicates polarised mitochondria and green indicates depolarised mitochondria. CCCP served as a positive control. Cells were counterstained with Hoechst and imaged on an EVOS M5000 imager. Corrected total cell fluorescence (CTCF) was used for quantification.

### 2.7 siRNA-mediated Drp-1 knockdown

Transient silencing of Drp-1 (encoded by DNM1L) in HCT-116 cells was carried out using a DNM1L-targeting small interfering RNA (siRNA) purchased from Santa Cruz Biotechnology (sc-43732). An in-house-produced non-targeting scrambled siRNA was used as the negative (Scrm) control. Cells were seeded in 6-well plates at 0.3 × 10 cells/well and grown to approximately 60–70% confluency. Transfection was performed at a final siRNA concentration of 20 nM using Lipofectamine RNAiMAX diluted in Opti-MEM™ reduced-serum medium (Gibco), following the manufacturer’s protocol. After 6 h, the transfection medium was replaced with complete growth medium. Knockdown efficiency was confirmed by qRT-PCR, and the expression of Drp-1, MID49 and MID51 was quantified as described in Section 2.8. All transfections were performed in three independent biological replicates.

### 2.8 Quantitative real-time PCR

Total RNA was extracted from control and HSC-CM-treated CRC cells after 48 h of treatment using TRIzol™, and quantified using a Colibri Titertek Nanophotometer (Berthold, Germany). One microgram of RNA was reverse-transcribed using iScript RT (Bio-Rad, 1708891), and qRT-PCR was performed using SsoFast™ EvaGreen Supermix (Bio-Rad, 1725201) per the manufacturer’s instructions (Extended File, Table 4). mRNA levels were normalised to GAPDH and relative fold-changes calculated using the 2^(–ΔΔCt) method (Extended File, Table 1).

### 2.9 Cellular and mitochondrial ROS measurement

After 48 h of HSC-CM treatment, HCT-116 cells were washed with PBS and stained with DCFDA (50 µM) or MitoSOX (500 nM) for 20–30 min. After washing, DCFDA-stained cells were imaged using the EVOS M5000 fluorescent imager, and relative intensity was calculated as (DCFDA intensity / Hoechst intensity) × 100. MitoSOX-stained cells were analysed by flow cytometry (Beckman Coulter) using FCS Express software.

### 2.10 Metabolic flux analysis

Oxygen consumption rate (OCR) was measured in HCT-116 cells using the Seahorse XFp Mitochondrial Stress Test (Agilent Technologies, USA). Cells were seeded at 1 × 10 cells/well in complete medium and incubated overnight at 37 °C with 5% CO. The next day, cells were treated with HSC-CM for 48 h, and OCR was measured using a Seahorse XFp analyser. The assay medium was supplemented with pyruvate, glutamine and glucose 1 h before reading. Injections were performed sequentially with oligomycin, FCCP and rotenone/antimycin A. Each experiment was performed in quadruplicate.

### 2.11 MitoTracker assay

To quantify polarised, actively respiring mitochondria, control and HSC-CM-treated HCT-116 cells were stained with MitoTracker Red CMXRos (400 nM) and Hoechst 33342 for 20 min. Because CMXRos accumulation depends on mitochondrial membrane potential, this readout reports the content of polarised mitochondria rather than total mitochondrial mass. Cells were imaged on an EVOS M5000 fluorescent imager and analysed using ImageJ. The relative content of polarised mitochondria was determined by comparing MitoTracker Red and Hoechst fluorescence intensities between control and treated cells; no inference regarding total mitochondrial mass was drawn from this measurement.

### 2.12 RNA sequencing

Total RNA from control and HSC-CM-treated CRC cells was isolated and quality-checked. Libraries were prepared using TruSeq® Stranded mRNA Library Prep (Illumina, USA) and sequenced on a NovaSeq 6000 platform using an S4 paired-end 200-cycle flow cell. Post-sequencing analysis was performed using the standard Illumina DRAGEN RNA-Seq pipeline. Pathway enrichment was performed using Enrichr; differentially expressed genes (DEGs) were clustered using Uniform Manifold Approximation and Projection (UMAP) and ranked by p-value using Reactome 2022. Gene ontology and KEGG enrichment was performed in Metascape, and mitochondrial DEGs were annotated using MitoCarta v3.0.

### 2.13 Mitochondrial fractionation and Western blot

Cells were washed with ice-cold PBS and lysed in mitochondrial lysis buffer (250 mM sucrose, 20 mM HEPES pH 7.4, 20 mM NaF, 10 mM KCl, 1.5 mM MgCl, 1 mM EDTA, 1 mM EGTA, 1 mM DTT, 200 mM benzamidine, 40 µg/mL leupeptin and 200 mM PMSF). After brief sonication, lysates were centrifuged at 1,000 × g for 10 min at 4 °C to remove debris and nuclei; the supernatant was further centrifuged at 13,000 × g for 20 min at 4 °C to separate cytosolic (supernatant) and mitochondrial (pellet) fractions. The mitochondrial pellet was washed twice with lysis buffer, resuspended in the same buffer, resolved by SDS-PAGE and analysed by Western blotting.

### 2.14 Statistical analysis

All experiments were performed as three independent biological replicates unless stated otherwise. Throughout, ‘biological replicate’ denotes an independent experiment performed on a separate passage of cells; imaging fields, Seahorse wells and flow-cytometry measurements within an experiment are technical replicates. The exact n and the experimental unit are stated in each figure legend. Statistical analyses were carried out using GraphPad Prism 8 (GraphPad Software, San Diego, USA), R (v4.2.2) and Microsoft Excel. Data are presented as mean ± SEM and mean +/− SD, and p < 0.05 was considered significant. Comparisons between two groups were made using Student’s t-test; comparisons among more than two groups were made using one-way ANOVA, as indicated in the figure legends.

## 3. Results

### 3.1 HSC-derived conditioned media alter mitochondrial metabolites in CRC cells

To obtain an overview of metabolic alterations in HSC-CM-treated HCT-116 cells, we performed DIA-based global metabolomic profiling. We identified 613 metabolites in MS1 positive mode and 541 metabolites in MS1 negative mode. At MS2 level, 136 metabolites were identified in negative mode and 320 in positive mode. Enrichment of dysregulated metabolites in biological pathways revealed prominent effects on the mitochondrial electron transport chain, mitochondrial dynamics, lipid metabolism (arachidonic acid metabolism, steroidogenesis, α-linolenic and linoleic acid metabolism, steroid biosynthesis and fatty acid biosynthesis) [11], suggesting that HSC-CM modulates inflammatory signalling and mitochondrial membrane dynamics in CRC cells (Fig. 1a–b and Supplementary Fig. 1).

**Fig. 1.**
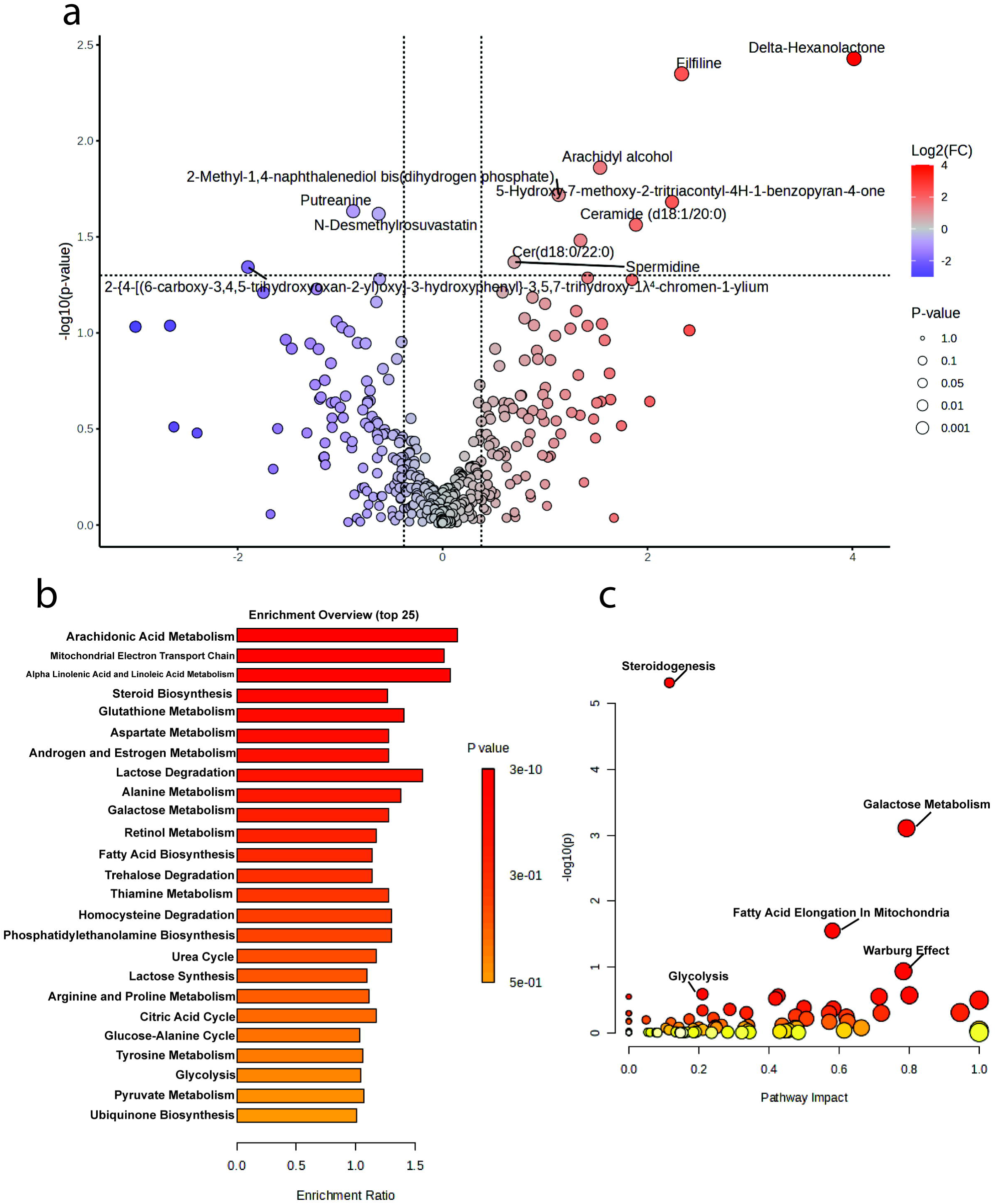
Metabolomic analysis of HSC-CM-treated CRC cells. (a) Volcano plot showing differentially expressed metabolites in HSC-CM-treated CRC cells; the x-axis represents log fold-change and the y-axis represents –log (p-value). (b) Enrichment analysis of metabolic pathways: the bar graph shows the top 25 enriched pathways, with bars colour-coded by enrichment p-value. (c) Scatter plot illustrating the impact of different metabolic pathways; the x-axis represents pathway impact score and the y-axis represents –log (p-value). Pathways of interest are labelled.

### 3.2 Conditioned media perturb mitochondrial function and related pathways

Next, we explored the impact of HSC-CM at the proteomic level. Proteomic analysis revealed significant alterations in mitochondrial proteins in HSC-CM-treated HCT-116 cells. Functional enrichment analysis highlighted changes in biological processes, cellular components and molecular functions, with notable enrichment of mitochondrion-related terms (Fig. 2a). Several proteins involved in mitochondrial dynamics most notably COX2 and COX4I1 were substantially downregulated; both play key roles in the electron transport chain and ATP production (Fig. 2c) [13]. NDUFB4, a component of the complex I respirasome that interacts with UQCRC1 (a subunit of complex III) to support respirasome integrity, was likewise downregulated. Pathway analysis further confirmed the dysregulation of respiratory electron transport, ATP synthesis, cellular stress responses and broader metabolic pathways, alongside protein localisation, amino acid metabolism and mitochondrial biogenesis, indicating that HSC-CM substantially remodels the mitochondrial proteome in CRC cells (Fig. 2d).

**Fig. 2.**
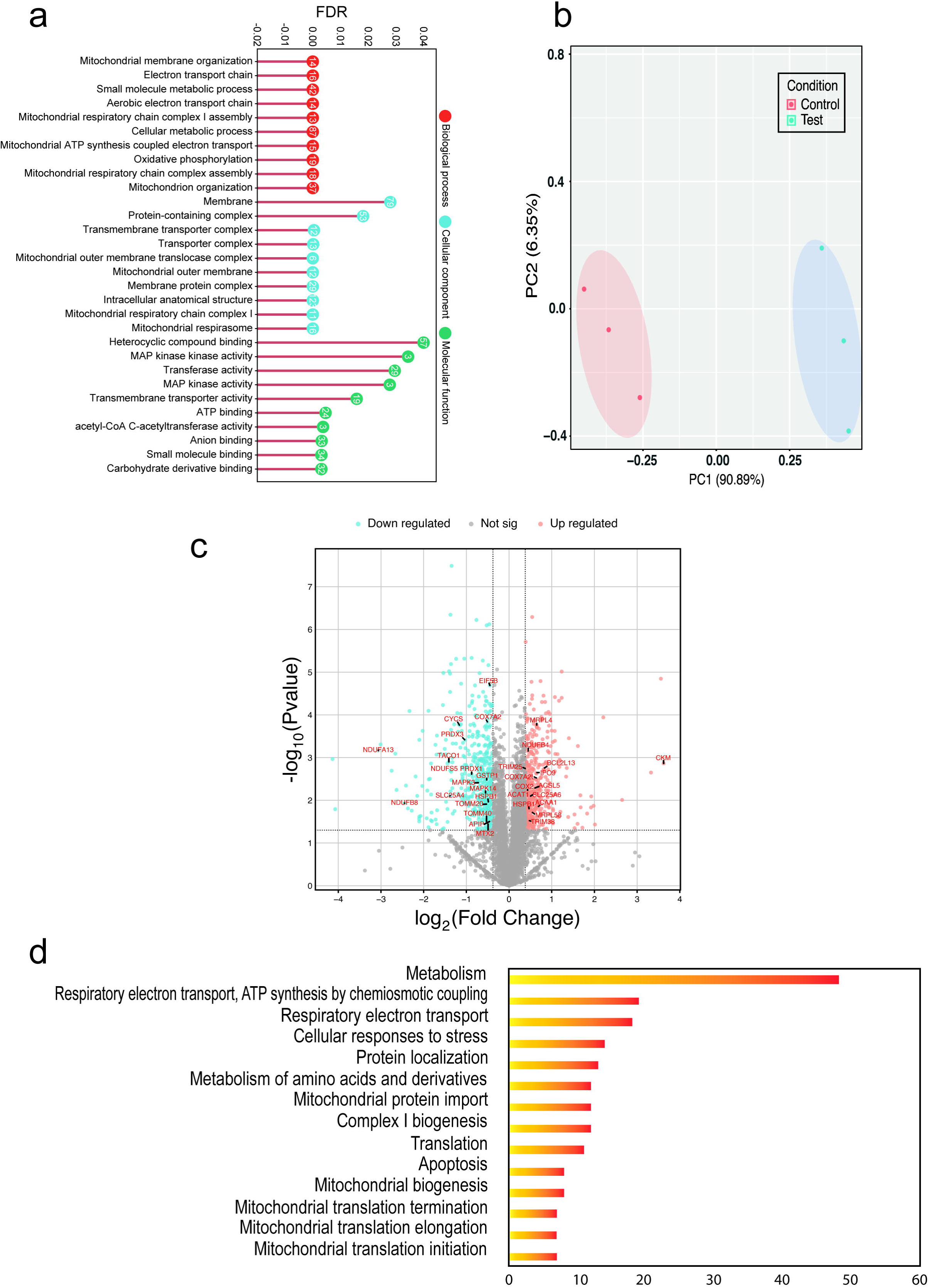
Proteomic analysis of HSC-CM-treated CRC cells. (a) Functional enrichment of differentially expressed proteins categorised by biological process, cellular component and molecular function. (b) Principal component analysis (PCA) of proteomic data, showing separation between control and HSC-CM-treated samples. (c) Volcano plot of differentially expressed proteins: red, significantly upregulated; blue, significantly downregulated; grey, non-significant changes. (d) Top enriched pathways based on differentially expressed proteins.

### 3.3 HSC-CM upregulates Drp-1 in CRC cells

Given that the proteomic data indicated profound alterations in mitochondrial dynamics, we next examined the fusion–fission machinery. Immunofluorescence analysis revealed only modest differences in OPA1 fluorescence intensity between control and treated cells (Fig. 3c–d); these panels quantify fluorescence intensity and not mitochondrial length. In contrast, Drp-1 expression was significantly increased in HSC-CM-treated cells compared with controls. Mitochondrial branch length, quantified from the mitochondrial morphology analysis, was markedly reduced and mitochondria appeared fragmented (p < 0.001) (Fig. 3a), with elevated Drp-1 fluorescence intensity (Fig. 3b). Moreover, we observed at the protein level the upregulation of both Drp 1 in phospho-serine 616 and 637, indicating that these sites are active. These findings indicate that HSC-CM drives excessive mitochondrial fission via Drp-1 upregulation.

**Fig. 3.**
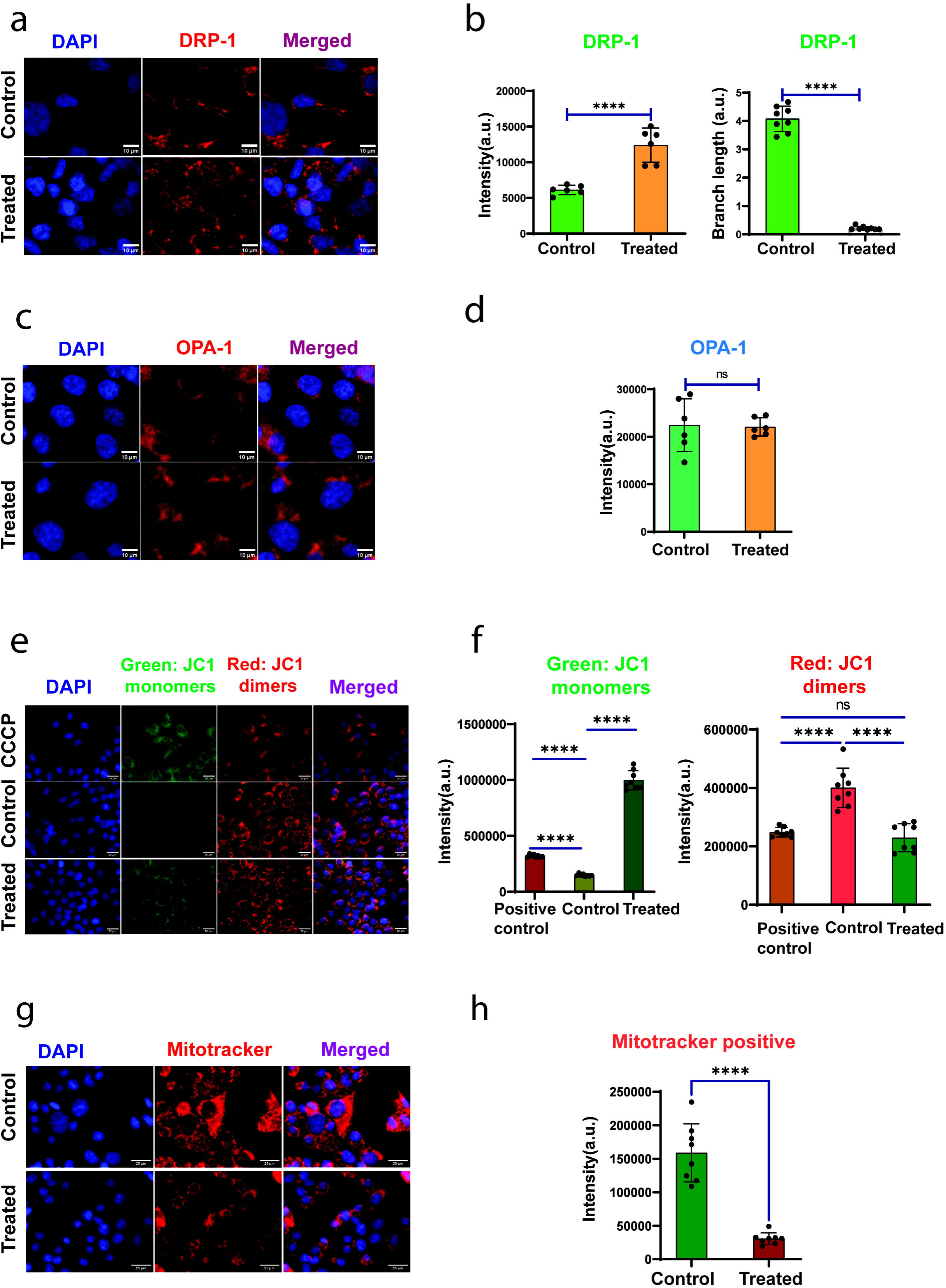
HSC-CM alters mitochondrial membrane potential and Drp-1 expression in CRC cells. (a) Immunofluorescence images of Drp-1 (DAPI, blue; Drp-1, red). Scale bar: 10 µm. (b) Quantification of Drp-1 intensity and mitochondrial branch length in control and HSC-CM-treated groups. (c) Immunofluorescence images of OPA1-stained CRC cells. (d) Quantification of OPA1 intensity in control and treated groups (n=6). (e) JC-1 fluorescence images of HSC-CM-treated CRC cells; scale bar: 20 µm. (f) Quantification of JC-1 green and JC-1 red intensities. (g) MitoTracker fluorescence images of HSC-CM-treated vs. control cells. (h) Quantification of active-mitochondria intensity, n=8. Scale bar: 20 µm. Data are mean ± SEM. Statistical significance was determined by one-way ANOVA (****p < 0.0001).

We then assessed mitochondrial membrane potential. HSC-CM treatment significantly increased JC-1 green-monomer accumulation (p < 0.0001) (Fig. 3e–f), reflecting membrane depolarisation and mitochondrial fragmentation comparable to that induced by the positive control CCCP. MitoTracker CMXRos staining further revealed a substantial reduction in polarised, actively respiring mitochondria in HSC-CM-treated cells (p < 0.0001) (Fig. 3g–h), consistent with the JC-1 depolarisation data; because CMXRos uptake is membrane-potential dependent, this was not interpreted as a change in total mitochondrial mass.

### 3.4 HSC-CM modulates the Drp-1 axis in CRC cells

To investigate the role of HSC-CM in mitochondrial fission, we performed siRNA-mediated Drp-1 knockdown (KD) and compared Drp-1 expression in knockdown and wild-type cells. Drp-1 knockdown significantly reduced Drp-1 expression (p < 0.0001), confirming efficient silencing, while HSC-CM treatment rescued Drp-1 levels in knockdown cells (KD + T; p < 0.001), indicating that HSC-CM directly drives Drp-1 upregulation. MID49 expression increased upon Drp-1 knockdown (p < 0.0001), potentially compensating for the loss of Drp-1, and was partially normalised by HSC-CM (p < 0.001). In contrast, MID51 remained suppressed in both KD and HSC-CM-treated groups (p < 0.0001), suggesting divergent regulatory roles for the MiD isoforms.

At the transcript level, mitochondrial fission genes Fis-1 (p < 0.001), Drp-1 (p < 0.001) and Parkin-2 (p < 0.0001) were upregulated in HSC-CM-treated cells compared with controls. Fusion-gene expression was largely unaffected (OPA1 p < 0.05; MFN1 and MFN2 not significant, p > 0.05) (Fig. 4b–c), consistent with selective induction of the fission programme.

**Fig. 4.**
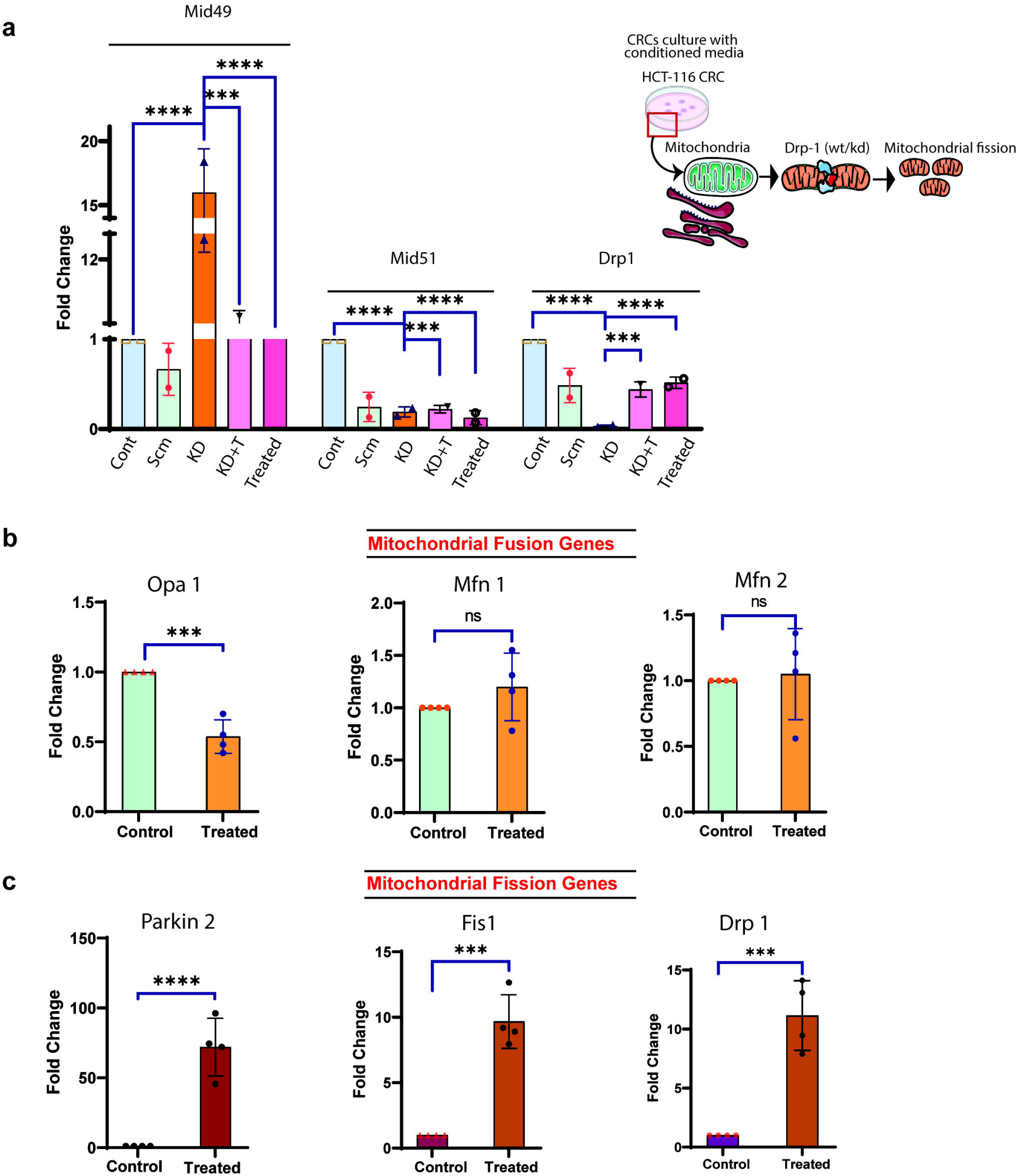
RT-qPCR analysis of mitochondrial fission and fusion gene expression in CRCs. (a) Expression of mitochondrial fission genes MID49, MID51 and DRP1 in CRC cells under different conditions: wild-type (Cont), scrambled control (Scrm), Drp-1 knockdown (KD), Drp-1 knockdown plus HSC-CM (KD + T) and HSC-CM treatment alone (Treated). Data are presented as fold change relative to control. The schematic illustrates the experimental design used to assess mitochondrial dynamics via Drp-1 modulation in HSC-CM-treated CRCs. (b) Expression of OPA1, MFN1 and MFN2 in control versus HSC-CM-treated cells. (c) Expression of Parkin-2, Fis-1 and Drp-1 in control versus HSC-CM-treated cells. Data are mean ± SEM. Statistical significance: ****p < 0.0001; ***p < 0.001; ns, not significant.

**Fig. 5.**
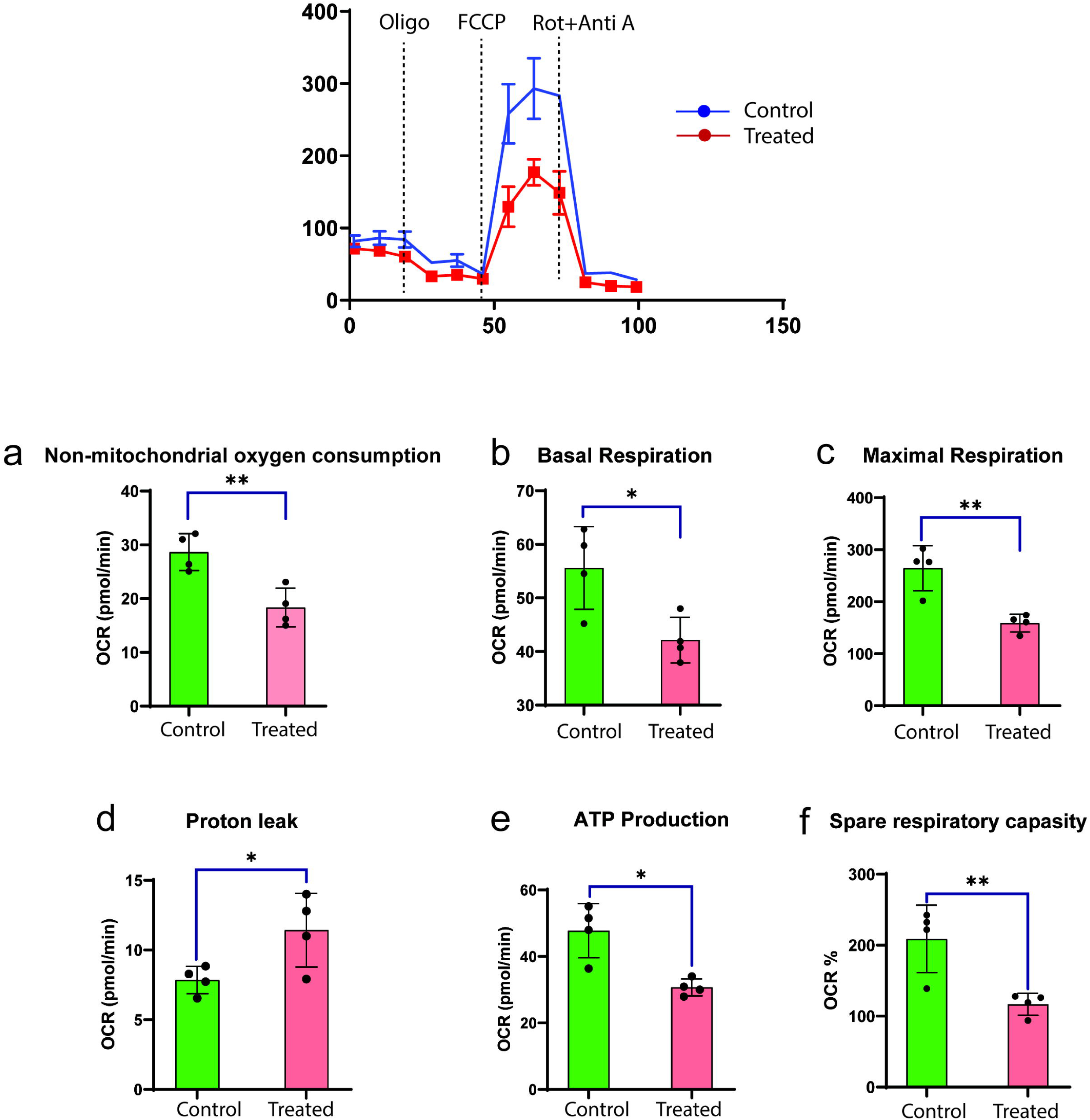
Effects of HSC-CM on mitochondrial respiration in CRC cells. Oxygen consumption rate (OCR) profiles of HCT-116 cells treated with HSC-CM versus control medium, measured using a Seahorse XF analyser. (a) Non-mitochondrial OCR decreased from ∼28.5 to ∼18.3 pmol/min (p < 0.01) (b) Basal respiration decreased by 24% (p < 0.05). (c) Maximal respiration was reduced by ∼40% (p < 0.01). (d) Proton leak increased 1.4-fold (p < 0.05), indicating uncoupling of oxidative phosphorylation. (e) ATP production declined by ∼36% (p < 0.05). (f) Spare respiratory capacity collapsed by ∼44% (p < 0.01). Data are mean ± SEM from n = 4 independent biological experiments; each biological experiment comprised four technical replicate wells on the Seahorse XFp cartridge. OCR values were normalised to post-assay cell number per well. Statistical significance was determined using an unpaired t-test.

### 3.5 HSC-CM-induced hyperfission drives mitochondrial dysfunction

Real-time metabolic flux was assessed in control and HSC-CM-treated HCT-116 cells using the Seahorse XFp system. HSC-CM induced severe mitochondrial dysfunction, evidenced by significant changes in OCR parameters. Basal respiration decreased by ∼24% (from ∼55.5 to 42 pmol/min; p < 0.05); maximal respiration and ATP production were each reduced by ∼40% (from ∼265 to 158 pmol/min & ∼36% from 47.5 to 30.5 respectively; p < 0.01 & p < 0.05); and spare respiratory capacity collapsed by ∼44% (from ∼208% to 117% of basal levels; p < 0.01). Proton leak increased 1.4-fold (from ∼7.9 to 11.4 pmol/min; p < 0.05), indicating uncoupling of oxidative phosphorylation. Collectively, these data indicate that HSC-derived secreted factors disrupt mitochondrial bioenergetics, impair respiratory capacity and precipitate an energy crisis in CRC cells [14].

### 3.6 HSC-CM induces mitophagy in treated CRC cells

We next measured total cellular and mitochondrial ROS in control and HSC-CM-treated cells. HSC-CM treatment markedly increased intracellular ROS (Fig. 6a–b) and the proportion of MitoSOX-positive cells increased from 0.37% in the stained control to 29.69% after HSC-CM treatment (positive control 18.21% ;Fig. 6c). Given the elevated ROS levels, we assessed apoptosis and observed a substantial increase in early (58.78%) and late (13.49%) apoptosis in treated cells (Fig. 6d). Western blotting confirmed upregulation of pro-apoptotic Bax and caspase-7 in HSC-CM-treated CRC cells (Fig. 6e–f & S3). Mitophagy markers were also examined: PINK1, the sensor of mitochondrial damage (Fig. 6f & S3), was significantly upregulated, and its partner Parkin showed increased expression in treated cells, indicating engagement of the PINK1–Parkin mitophagy pathway. These static measurements demonstrate activation of the pathway but do not by themselves quantify mitophagic flux; assays such as LC3-II turnover with and without lysosomal inhibition, mitochondria–lysosome colocalisation or a validated mitophagy reporter would be required to establish flux.

**Fig. 6.**
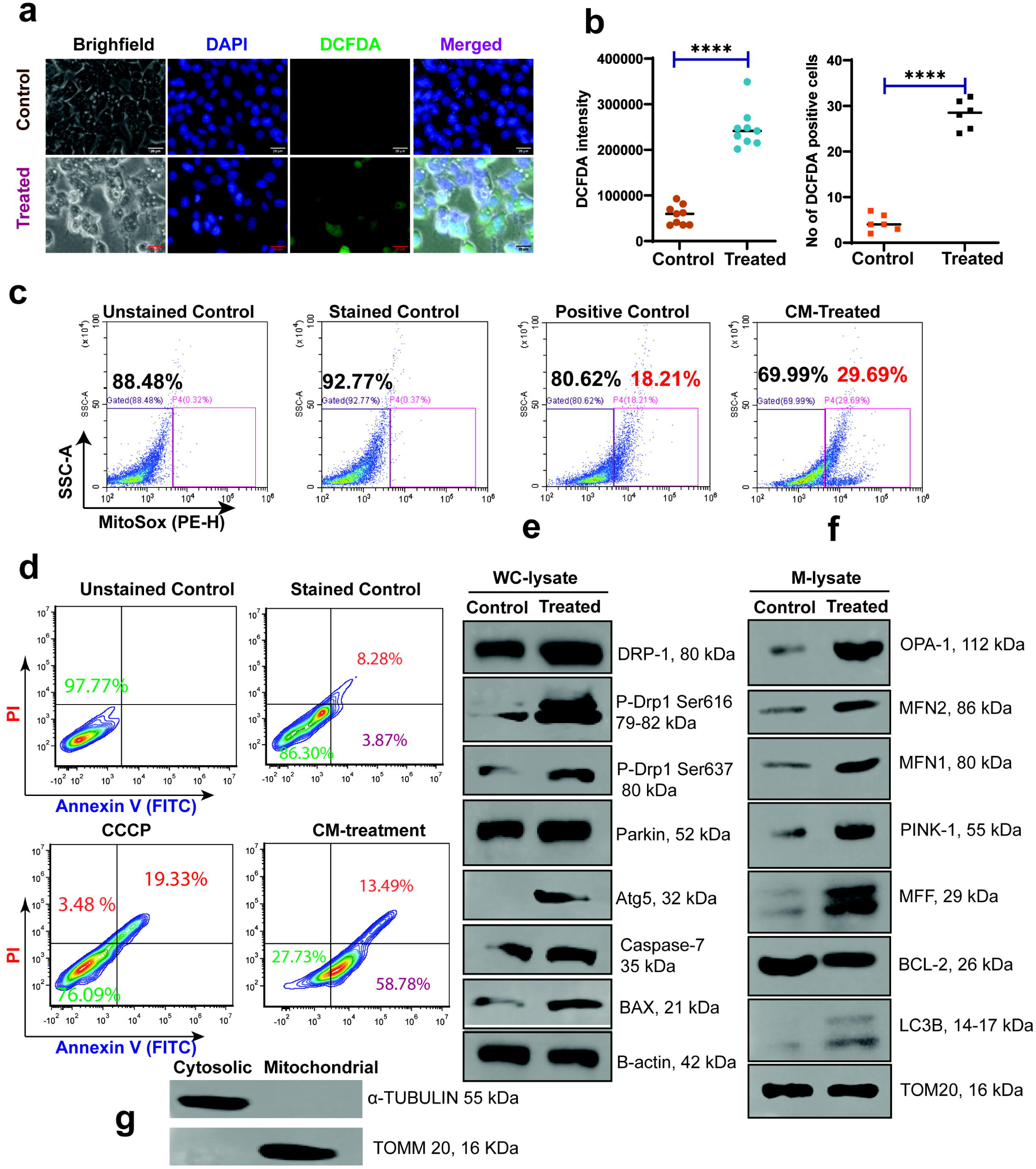
HSC-CM treatment alters redox state, induces mitochondrial dysfunction, and promotes apoptosis in CRC cells. (a) Representative DCFDA images showing intracellular ROS in CRC cells treated with HSC-CM versus control. Cells were stained with DAPI (blue), DCFDA (green) and the merged channel is shown. Scale bar 20 µm. (b) Quantification of DCFDA fluorescence intensity (left) and number of DCFDA-positive cells (right). Data are mean ± SD (n = 8); ****p < 0.0001 (unpaired t-test). (c) MitoSOX Red flow-cytometry plots showing mitochondrial superoxide levels; positive control and HSC-CM-treated cells display significantly increased MitoSOX positivity. (d) Annexin V/PI flow cytometry showing cell viability and apoptosis. (e–f) (e) whole-cell lysate, control and treated, normalised to beta-actin; panel (f) mitochondrial fraction, control and treated, normalised to TOM20; panel (g) fraction-purity control.

### 3.7 RNA-Seq reveals HSC-CM-induced mitophagy and autophagy in treated CRCs

To further evaluate apoptosis and autophagy-mediated mitochondrial clearance, RNA-Seq was performed on control and HSC-CM-treated HCT-116 cells. More than 126 genes were upregulated and six genes were downregulated in treated cells (Fig. 7b–c). Many of the upregulated genes including ATG5, VMP1, BCLAF1, XIAP, MSH2 and KLF6 are closely associated with mitochondrial fission, mitophagy and apoptosis (Fig. 7a). Increased ATG5 protein expression was independently confirmed by Western blotting (Fig. 6f). Excessive mitochondrial division is a known driver of mitophagy (via ATG5 and VMP1) and sensitises cells to apoptotic signals (via BCLAF1 and XIAP), particularly under stress [15–17]. Activation of stress-responsive transcription factors (KLF6, NFE2L3), DNA-repair and stress-response genes (MSH2, POLI, DNAJA1) [18], and the concurrent upregulation of mitochondrial-protein components (MRPS33), protein-degradation systems (PSMA3, PSMC1) and RNA helicases (DDX5, DDX6, DDX18) collectively indicate a coordinated response geared toward removal of damaged mitochondria, induction of apoptosis in persistently damaged cells, and resolution of cellular stress. Gene Ontology analysis confirmed enrichment of dysregulated genes in mitochondrion-protein complexes, inner-membrane components and ribosomal biogenesis (Fig. 7d–f).

**Fig. 7.**
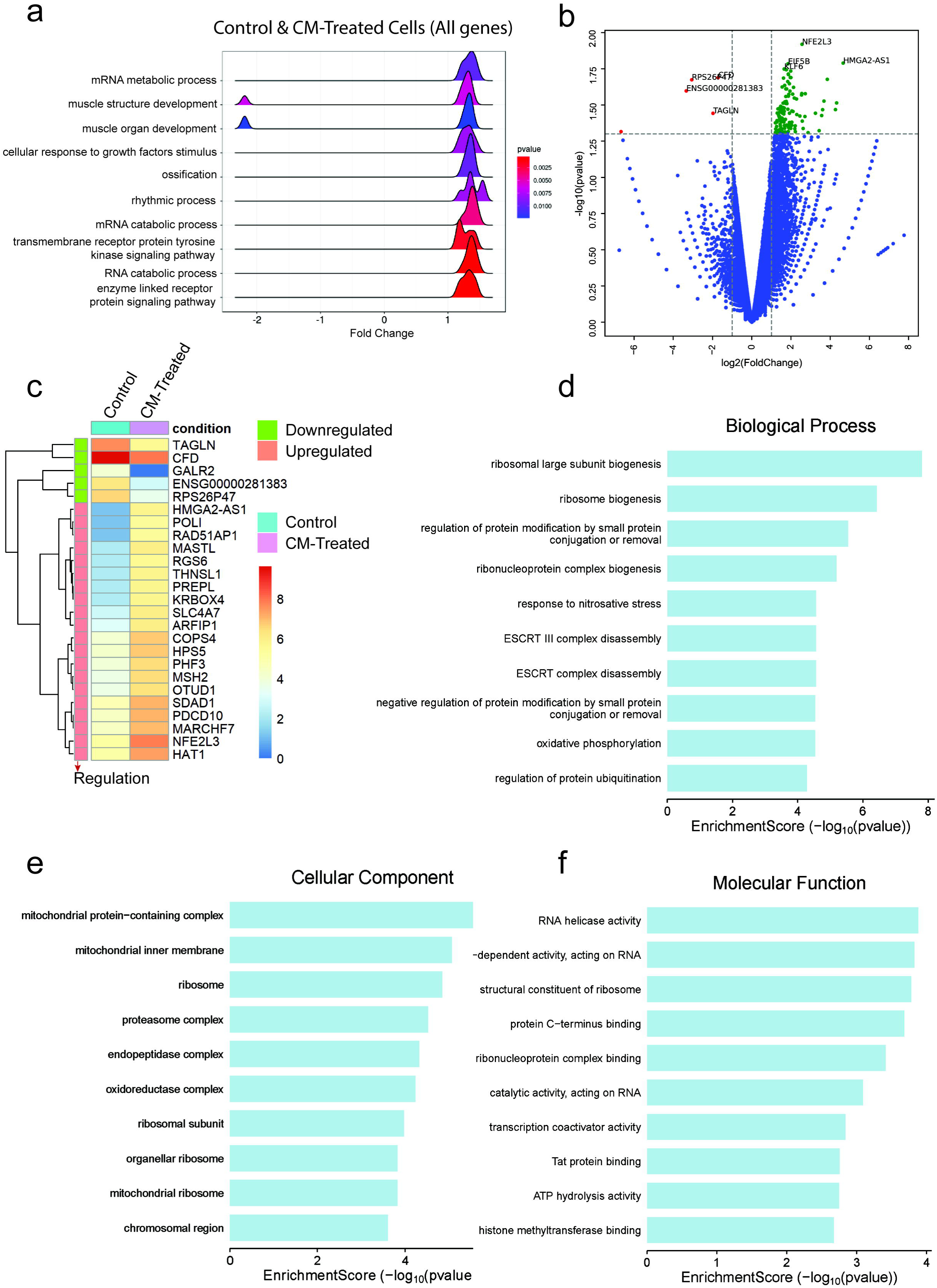
Transcriptomic profiling of HSC-CM-treated CRC cells. (a) Ridge plot showing gene-set enrichment for key biological processes between control and HSC-CM-treated cells. (b) Volcano plot showing significantly up- and downregulated genes upon HSC-CM treatment. (c) Heatmap of top differentially expressed genes between groups. (d–f) GO enrichment analyses revealing affected biological processes (d), cellular components (e) and molecular functions (f), with particular emphasis on ribosome biogenesis, mitochondrial complexes and RNA metabolism. Cut-off: p ≤ 0.05 and |log FC| ≥ 1.

## 4. Discussion

Hematopoietic stem cells (HSCs) are indispensable for sustaining the blood-cell compartment and contributing to tissue repair, but their paracrine and secretory influences on solid tumours, particularly CRC, remain largely unexplored. Our previous work [11] showed that HSC-CM contain a range of cytokines that modulate mitochondrial bioenergetics; the present study demonstrates, for the first time, that HSC-CM exerts a potent anti-tumour effect on CRC cells by specifically targeting mitochondrial dynamics a master regulator at the intersection of cancer bioenergetics, metabolic plasticity and survival [18–20].

Mitochondrial dynamics, governed by a delicate balance between fusion and fission, orchestrate mitochondrial morphology, quality control and function [21]. Dysregulation of this balance is increasingly recognised as a hallmark of cancer [22]. Our data show that HSC-CM skews this equilibrium decisively toward excessive fission through the upregulation of Drp-1 and Fis-1, while the fusion proteins MFN1, MFN2 and OPA1 didn’t show any major alteration in their expression. This selective induction triggers widespread mitochondrial fragmentation and disintegration of the mitochondrial network.

The consequences of this hyper-fragmented state are profoundly detrimental to cancer-cell viability [23]. Excessive fission precipitates depolarisation of the mitochondrial membrane potential (ΔΨm) (Fig. 3), an early hallmark of mitochondrial distress that disrupts the proton gradient required for ATP synthesis [24,25]. Concurrent disruption of the electron transport chain evidenced by downregulation of COX2 (complex IV) and NDUFB4 (complex I) results in diminished OXPHOS capacity and a bioenergetic crisis with significantly reduced ATP output. Seahorse XF analyses corroborated this collapse, showing decreased basal and maximal respiration alongside increased proton leak.

We further examined whether HSC-CM activity translates to a more physiologically relevant 3D context. In HCT-116 spheroid cultures, HSC-CM treatment produced marked reductions in spheroid size and density relative to controls, with significant decreases in spheroid area and cellularity (Supplementary Fig. 2a–b). This disruption of 3D tumour architecture is a direct indicator of compromised tumour growth and self-renewal in a physiologically relevant environment [26–28]. Viability was further assessed by MTT, which confirmed a strong anti-proliferative effect: HSC-CM-treated CRC cells showed a more than three-fold decrease in proliferation compared with controls (Supplementary Fig. 2d; p < 0.001). These data support the conclusion that HSC-derived secretory factors contribute to metabolic and proliferative collapse within the tumour microenvironment. The metabolic reprogramming induced by mitochondrial dysfunction is also reflected in our FTIR spectra (Supplementary Fig. 2c): increased transmittance in HSC-CM-treated CRC supernatants indicates a reduction in IR-absorbing metabolic by-products, consistent with impaired mitochondrial activity and decreased secretion of metabolites such as TCA-cycle intermediates and amino-acid catabolites. This altered extracellular metabolite signature highlights how mitochondrial integrity governs cellular metabolic output [29–31].

Mitochondrial dysfunction also elevates intracellular ROS [32], which we confirmed directly via increased DCFDA and MitoSOX staining (Fig. 6). Elevated mitochondrial superoxide can act as a second messenger to activate intrinsic apoptotic pathways [33,34]. Consistent with this, increased expression of Bax and caspase-7 indicates initiation of the intrinsic apoptotic programme, driving programmed cell death. This oxidative stress-driven apoptosis is a key mechanism by which HSC-CM mediates CRC-cell lethality.

To counter the accumulation of damaged mitochondria, cells activate mitochondrial quality-control mechanisms. Our data show clear activation of the PINK1–Parkin mitophagy pathway in response to HSC-CM, as evidenced by increased PINK1 and Parkin protein expression (Figs 4 and 6), alongside upregulation of mitophagy/autophagy markers including ATG5 (Figs 6f and 7). This mitophagy response represents an attempted salvage mechanism; however, given the magnitude of mitochondrial damage and bioenergetic failure, it is unable to restore homeostasis and instead coincides with increased apoptosis.

Mechanistically, Drp-1 siRNA knockdown experiments revealed that HSC-CM-derived secretory factors can rescue Drp-1 expression in silenced cells, underscoring their direct role in driving mitochondrial fission [35,36]. This points to a previously unrecognised extracellular regulatory axis through which the HSC secretome modulates mitochondrial morphology and function in distal cancer cells.

Integration of our multi-omics data transcriptomics, metabolomics and proteomics further supports a model in which HSC-CM induces a coordinated cellular stress response centred on mitochondrial dysfunction but also affecting oncogenic signalling pathways such as mTOR and MYC (Supplementary Fig. 1), as well as autophagy and cellular clearance machinery. This comprehensive remodelling culminates in CRC clearance, positioning mitochondrial dynamics not only as a vulnerability but also as a therapeutically exploitable target.

Previous studies on mitochondrial fission in cancer have underscored a dichotomy: moderate fission supports tumour progression, metastasis and chemoresistance by enabling rapid mitochondrial redistribution and metabolic adaptation, whereas excessive fission triggers bioenergetic failure and apoptosis. Our work positions HSC-CM as a novel, biologically derived modulator capable of pushing CRC cells beyond this threshold, tipping mitochondrial fission toward a lethal hyper-fragmented state [17,19,34–36].

Collectively, these findings highlight the mitochondrial fission machinery and its quality-control pathways as promising non-toxic therapeutic targets. Exploitation of the HSC secretome or of its active components provides an innovative adjunct or alternative to conventional chemotherapy, with potential to overcome resistance mechanisms rooted in mitochondrial and metabolic plasticity [2,6,37].

## 5. Conclusion

Our results demonstrate that manipulation of mitochondrial dynamics by the HSC secretome drives CRC cells beyond their metabolic tipping point, disrupting bioenergetic homeostasis, inducing oxidative stress-mediated apoptosis and activating mitophagy. This study lays the groundwork for novel mitochondrial and metabolic interventions in colorectal cancer, leveraging endogenous stem-cell factors to selectively impair the survival of tumour cells.

## Statements and Declarations

### Ethics approval

No ethics approval was required for this study. The work was conducted exclusively using commercially available, authenticated human cell lines (HCT-116 and HL-60) obtained from the National Centre for Cell Science (NCCS), Pune, India. The study did not involve human participants, identifiable human material, identifiable data, or animal experiments.

### Consent to participate

Not applicable. This study did not involve human participants.

### Consent to publish

Not applicable. The manuscript does not contain any individual person’s data in any form.

### Availability of data and materials

The raw metabolomics data generated during this study are publicly available in the Zenodo repository (DOI: https://doi.org/10.5281/zenodo.16736809; direct access: https://zenodo.org/records/16736809). Mass spectrometry–based proteomics data have been deposited in the ProteomeXchange Consortium via the PRIDE partner repository under dataset identifier **PXD066198**, accessible at https://www.ebi.ac.uk/pride/archive/projects/PXD066198. Reviewer can access the dataset by logging in to the PRIDE website using the following account details:Username, Password: tQ6UXt4Ekluc. RNA sequencing data have been submitted to the NCBI BioSample database, an INSDC member repository (SRA: SRS29407072); the accession number is PRJNA1474307. All other data generated or analysed during the current study are included in this published article and its supplementary information files, or are available from the corresponding authors upon reasonable request.

### Authors’ contributions (CRediT)

S.M.: Conceptualization, Investigation, Formal analysis, Methodology, Writing – original draft. A.B.R. and V.K.N.: Investigation (proteomics), Formal analysis. T.S.K.P.: Supervision (proteomics), Resources, Formal analysis. S.B.: Conceptualization, Funding acquisition, Writing – review & editing. S.S.P.: Supervision, Writing – review & editing. B.B.: Conceptualization, Supervision, Project administration, Writing – review & editing. All authors read and approved the final manuscript.

## Acknowledgements

The CSBMM authors acknowledge the support of the Department of Biotechnology, Government of India, to Yenepoya (Deemed to be University) through the project on “Skill Development in Mass Spectrometry-based Metabolomics Technology BIC” (BT/PR40202/BTIS/137/53/2023), and the Karnataka Biotechnology and Information Technology Services (KBITS), Government of Karnataka, for infrastructural support of the CSBMM at Yenepoya (Deemed to be University) under the Biotechnology Skill Enhancement Program in Multiomics Technology (BiSEP GO ITD 02 MDA 2017). S.M., A.B.R. and V.K.N. acknowledge fellowships from Yenepoya (Deemed to be University). We thank Dr. Shyam Prasad Varija Raghu (Associate Professor, Yenepoya Research Centre) for assistance with confocal image acquisition; Dr. Pavan S. R. (Post-Doctoral Research Associate, Department of Cellular and Molecular Biology, The University of Texas at Tyler School of Medicine) for help with metabolomics data analysis; and Ms. Anwesha Bose (CSIR-Senior Research Fellow, CSIR-Indian Institute of Chemical Biology, Kolkata) for assistance with Seahorse analysis. The authors also thank the Yenepoya Research Centre, Yenepoya (Deemed to be University), Mangalore, for infrastructural and administrative support.

## Clinical trial number

Not applicable.

## Supplementary

**Supplementary Fig. 1.** (a) Chord diagram visualising the relationship between proteins and interacted metabolites in HSC-CM-treated CRC cells. (b) Gene Ontology analysis of protein targets of HSC-CM treatment, highlighting enriched biological processes and the correlation of dysregulated metabolites with cellular component, biological process and molecular function categories.

**Supplementary Fig. 2.** (a–b) Spheroid culture assay of HCT-116 cells, and (b) quantification of spheroid area and intensity after HSC-CM treatment. (c) FTIR spectra of HCT-116 and HSC-CM-treated HCT-116 supernatants. (d) MTT assay of media+growth factors, 100% complete conditioned media, and 50-50% conditioned media-based HCT-116 cells after 48 h. Data are mean ± SEM; statistical significance was determined by one-way ANOVA (****p < 0.0001).

**Supplementary Fig. 3.** Quantifications of western blots of both whole cell lysate and mitochondrial lysate of HCT-116.

## Abbreviations

CCCP: carbonyl cyanide m-chlorophenylhydrazone
CD: cluster of differentiation
CM: conditioned media
CRC: colorectal cancer
DMSO: dimethyl sulfoxide
Drp-1: dynamin-related protein 1
ETC: electron transport chain
FACS: fluorescence-activated cell sorting
HSCs: hematopoietic stem cells
mCRC: metastatic colorectal cancer
MMP: mitochondrial membrane potential
MTT: 3-(4,5-dimethylthiazol-2-yl)-2,5-diphenyltetrazolium bromide
OCR: oxygen consumption rate
OXPHOS: oxidative phosphorylation
PBS: phosphate-buffered saline
PVDF: polyvinylidene fluoride
ROS: reactive oxygen species
WT: wild type

